# Identification of key candidate genes and pathways in axial Spondyloarthritis through integrated bioinformatics analysis

**DOI:** 10.1101/2020.03.17.995134

**Authors:** Zhen-zhen Zhang, Jing Zeng, Hai-hong Li, Yu-cong Zou, Shuang Liang, Gang Liu

**Affiliations:** Department of Rehabilitation medicine, The Third Affiliated Hospital of Southern Medical University, Guangzhou 510000, China; Department of Orthopedics, Tongji Hospital, Tongji Medical College, Huazhong University of Science and Technology, Wuhan 430030, China; Department of Oral Medicine, Infection and Immunity, Harvard School of Dental Medicine, Boston, MA 02115, USA

**Author notes:** Correspondence should be addressed to Prof. Gang Liu and Shuang Liang,;, Postal address: 183# West Zhongshan Avenue, Tianhe District, Guangzhou, 510000, China, Department of Orthopedics, Tongji Hospital, Tongji Medical College, Huazhong University of Science and Technology, Wuhan 430030, China.

**Keywords:** Axial Spondyloarthritis, differentially expressed genes, bioinformatics analysis, pathomechanisms

## Abstract

**Background:** Radiographic axial Spondyloarthritis (r-axSpA) is the prototypic form of seronegative spondyloarthritis (SpA). In the present study, we evaluated the key genes related with r-axSpA, and then elucidated the possible molecular mechanisms of r-axSpA.

**Material/Methods:** The gene expression GSE13782 was downloaded from the GEO database contained five proteoglycan-induced spondylitis mice and three naïve controls. The differentially expressed genes (DEGs) were identified with the Bioconductor affy package in R. Gene Ontology (GO) enrichment and the Kyoto Encyclopedia of Genes and Genomes (KEGG) pathway analysis were built with the DAVID program followed by construction of a protein-protein interaction (PPI) network performed with Cytoscape. WebGestalt was performed to construct transcriptional regulatory network and microRNAs-target regulatory networks. RT-PCR and immunohistochemical staining were performed to testify the expression of hub genes, transcription factors (TFs) and microRNAs.

**Results:** A total of 230 DEGs were identified. PPI networks were constructed by mapping DEGs into STRING, in which 20 hub proteins were identified. KEGG pathway analyses revealed that the chemokine, NOD-like receptor, IL-17, and TNF signalling pathways were altered. GO analyses revealed that DEGs were extensively involved in the regulation of cytokine production, the immune response, the external side of the plasma membrane, and G-protein coupled chemoattractant receptor activity. The results of RT-PCR and immunohistochemical staining demonstrated that the expression of DEGs, TFs and microRNAs in our experiment were basically consistent with the predictions.

**Conclusions:** The results of this study offer insight into the pathomechanisms of r-axSpA and provide potential research directions.

## 1. Introduction

Ankylosing spondylitis (axSpA) is a chronic autoimmune disease that involves the axial skeleton (1). AxSpA with spine(2) attachment point inflammation ultimately develops into pathological ossification and rigidity, which is the main cause of disability and the declining quality of life in patients with axSpA. Patients with ankylosing spondylitis have inflammatory pain and stiffness in the joints and spine, severely affecting range of activity. As a complicated chronic autoimmune inflammatory disease, in addition to invading the sacroiliac joint, spine, and spine soft tissue, axSpA can also cause extra-joint manifestations such as anterior uveitis and gut inflammation (3). Its pathological mechanism may be related to an imbalance of intestinal flora. Recent studies have confirmed that, to a certain degree, dysbiosis of intestinal flora participates in the pathogenesis of axSpA (4). The main clinical symptoms include pain and stiffness of the waist and hip, and joint swelling and pain in the limbs. AxSpA pain usually occurs as insidious-onset lower back and hip pain, and the condition worsens at night. Pain correlates with activation of the autoimmune system and neural pain signal, which are defined as neuro-immune interactions(5).

The influencing factors of axSpA such as gut dysbiosis and the *HLA-B27* gene, all primarily participate in its pathological processes (6). The *HLA-B27* gene is relevant to the pathogenesis of axSpA. It physiologically completes the folding and modification of peptide chain molecules in the endoplasmic reticulum, and its misfolding leads to the occurrence of axSpA (7). *HLA-B27* is a histocompatibility complex class I molecule expressed in all nucleated cells. *HLA-B27* homomorphic bivalent bind to receptors expressed on natural killer cells, granulocytes, and lymphocytes, thereby initiating a series of downstream signalling and participating in a series of cellular regulation processes (8). Therefore, it is bound to play an important role in the pathogenesis of autoimmune process. Studies have confirmed the high therapeutic effect of TNF-α blockers (golimumab, adalimumab, and etanercept) in the prevention and arrest of onset in patients with axSpA (4), although their prevention ability remains controversial. About 60% of axSpA patients respond to antibiotics (9). Numerous studies have verified the curative effects of non-steroid anti-inflammatory drugs (NSAIDs) on axSpA, although some may increase the risk of developing stomach ulcers (10). NSAIDs are most often used for the treatment of inflammatory pain symptoms because they block Cox enzymes and prostanoids (11). Recently, mesenchymal stem cells were utilised to relieve inflammatory responses and promote tissue regeneration through cell–cell interactions and release of cytokines, which is a novel therapeutic direction (12).

In this study, we obtained the candidate DEGs with R language. And then Kyoto Encyclopedia of Genes and Genomes (KEGG) pathway enrichment and PPI network were analysed and visualized. Subsequently, transcription factor (TF)‐mRNA regulatory network and miRNA-target genes regulatory network were constructed to identify the key regulatory genes. In conclusion, a total of 230 DEGs, 20 hub genes, 10 miRNAs and 10 TFs were identified.

## 2. Methods

### 2.1. Microarraydata

Gene expression profile data (GSE13782) derived from five proteoglycan-induced spondylitis (PGISp) mice (at weeks 0, 3, and 6) and three naïve controls were obtained from the GEO Database (www.ncbi.nlm.nih.gov/geo/query/acc.cgi?acc=GSE13782). The raw data (GSE13782) of high-throughput gene expression profiles were submitted by Laszlo A in 2019 and performed with GPL1261 platform. (Affymetrix Mouse Genome 430 2.0 Array).

### 2.2. Data pre-processing and identification of DEGs

The raw expression data (Affymetrix CEL files) were first pre-processed into expression values at the gene symbol level. Quantile normalisation was performed with the Robust Multiarray Average normalisation approach of the Bioconductor affy package and the preprocessCore package in R (13, 14). Finally, the limma package with multiple testing corrections was used to identify significant DEGs among the five PGISp mice and three naïve controls. The classic paired *t*-test was used to screen the statistically significant DEGs, and p < 0.05 and absolute value of log-fold change |logFC| > 1.5 were set as cut-off thresholds.

### 2.3. Gene ontology and pathway enrichment analyses of DEGs

To analyse the potential functions of candidate DEGs, GO functional enrichment analyses and KEGG pathway enrichment analyses were carried out with DAVID v6 .8, a tool for Annotation, Visualisation and Integrated Discovery (https://david.ncifcrf.gov/home.jsp). The DAVID database was used to analyse the functional relationship between GO and candidate DEGs (15). In our study, FDR ≤ 0.05 was considered the cut-off threshold for enriched annotation.

### 2.4. PPI networks construction and hub gene analyses

The STRING search tool was performed to build the PPI network (http://string-db.org/) (16). The DEG list was uploaded to STRING with a interaction score > ≥ 0.4 to build the PPI network. We analysed the numbers of DEGs between PGISp mice and blank control mice and further visualised the PPI network using the Cytoscape software. The nodes, edges of the network and the hub protein, were performed with the cytoHubba plugin (MCODE) with the criteria as follows: degree cutoff = 2, node score cutoff = 0.2, max Depth = 100 and k score=2. (17).

### 2.5. Construction and analyses of the TFs network

The WebGestalt (http://www.webgestalt.org/option.php) is an online open-source platform which utilizes established prediction platform and databases (18). Utilizing WebGestalt, we screened out significant TFs and correlative targeted genes. The Cytoscape software was used to build the DEG-associated transcriptional regulatory network. In this TF-targeted gene network, an adjusted P < 0.05 was considered the statistical standard for sorting out the most significant TF-targeted genes.

### 2.6. Construction and analysis of microRNAs-targeted gene network

WebGestalt was also used to screen distinctly different microRNAs and predict their target regulatory genes among the DEGs, in order to construct an microRNAs-targeted regulatory network. Finally, an microRNAs-targeted regulatory gene network was built and clearly visualised with the Cytoscape tool.

### 2.7. Mouse model, immunization and Histological Staining

The mouse was induced by injecting 2 month-old female BALB/c mice with2 mg human-derived proteoglycan (PG) together with 2 mg DDA (Sigma, MO, united States). as described previously (1). The PGISp models were injected 3 times at week 0, 3 and 6 of the study. All animal experiments were performed with the approval of the Ethical Committee of Tongji Hospital. Sacroiliac joint samples were collected 3 months after the first PG injection. After the mouses were sacrificed,, sacroiliac joint samples of per group were fixed in 4 % PFA for 24 hours followed by decalcification in 14 % EDTA for 2 weeks. Sacroiliac joint samples were cut into 4-μm serial sections along the coronal plane. Ten consecutive sections out of 30 were selected for IHC and HE staining. Antibodies against c-JUN (#9165s), c-FOS (#2250s) and STAT3 (#9139) were obtained from Cell Signaling Technology (Danvers, MA).

### 2.8. RNA extraction and RT-PCR analysis

Sacroiliac joint samples were homogenized and stored at −20°C. The total RNA kit (Toyobo, Japan) was used for RNA extraction. The Revert Aid First-Strand cDNA Synthesis Kit (Primescript RT, Takara) was used for the synthesis of Complementary DNA (cDNA). SYBR Green-based quantitative PCR were performed with the SYBR Premix Ex TaqTMII (Toyobo, Japan) with the specific primers (Supplementary Table 2).

### 2.9. Statistics

All experiments were repeated at least twice. Data are presented as mean±standard deviation. Statistical analyses were performed using Student’s t-test. p<0.05 was considered to be significant.

## 3. Result

### 3.1. Identification of DEGs

The GSE13782 profiles of PGISp mice and normal mice were analysed using the limma package. After data processing, a total of 230 DEGs were obtained, among which 36 were downregulated and 194 were upregulated (Supplementary Table 1). A heatmap of the differentially expressed genes is shown in Figure 1.

**Fig. 1.**
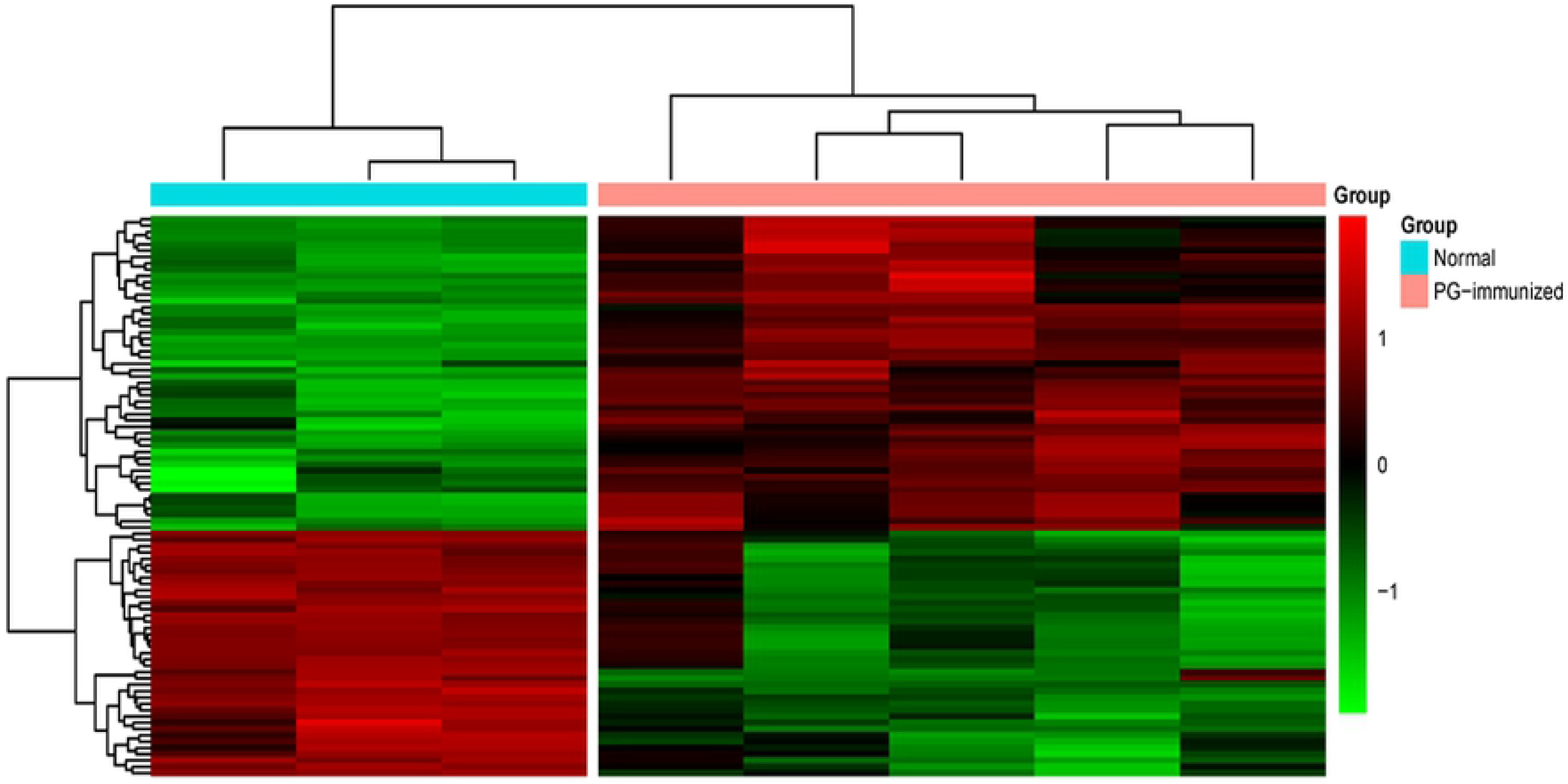
Heatmap plot of significantly DEGs identified in mice with ankylosing spondylitis.Red represents increased gene expression and green represents decreased gene expression.

### 3.2. GO and pathway enrichment analyses

To gain further understanding of the biological functions of the differentially expressed genes, the upregulated and downregulated genes were performed with the DAVID tool; the top 15 KEGG pathways are showed in Figure 2. The DEGs were widely involved in the chemokine, NOD-like receptor, IL-17, and TNF signalling pathways. GO function enrichment analyses revealed that 230 DEGs were enriched in 11 GO functions (FDR< 0.05; Fig. 3.), including 5 Biological Process (BP) terms, 1 Cellular Component (CC) term, and 5 Molecular Function (MF) terms. As shown in Figure 3, the DEGs were mainly involved in regulation of cytokine production, the immune response-regulating signalling pathway, the external side of the plasma membrane, and G-protein coupled chemoattractant receptor activity.

**Fig. 2.**
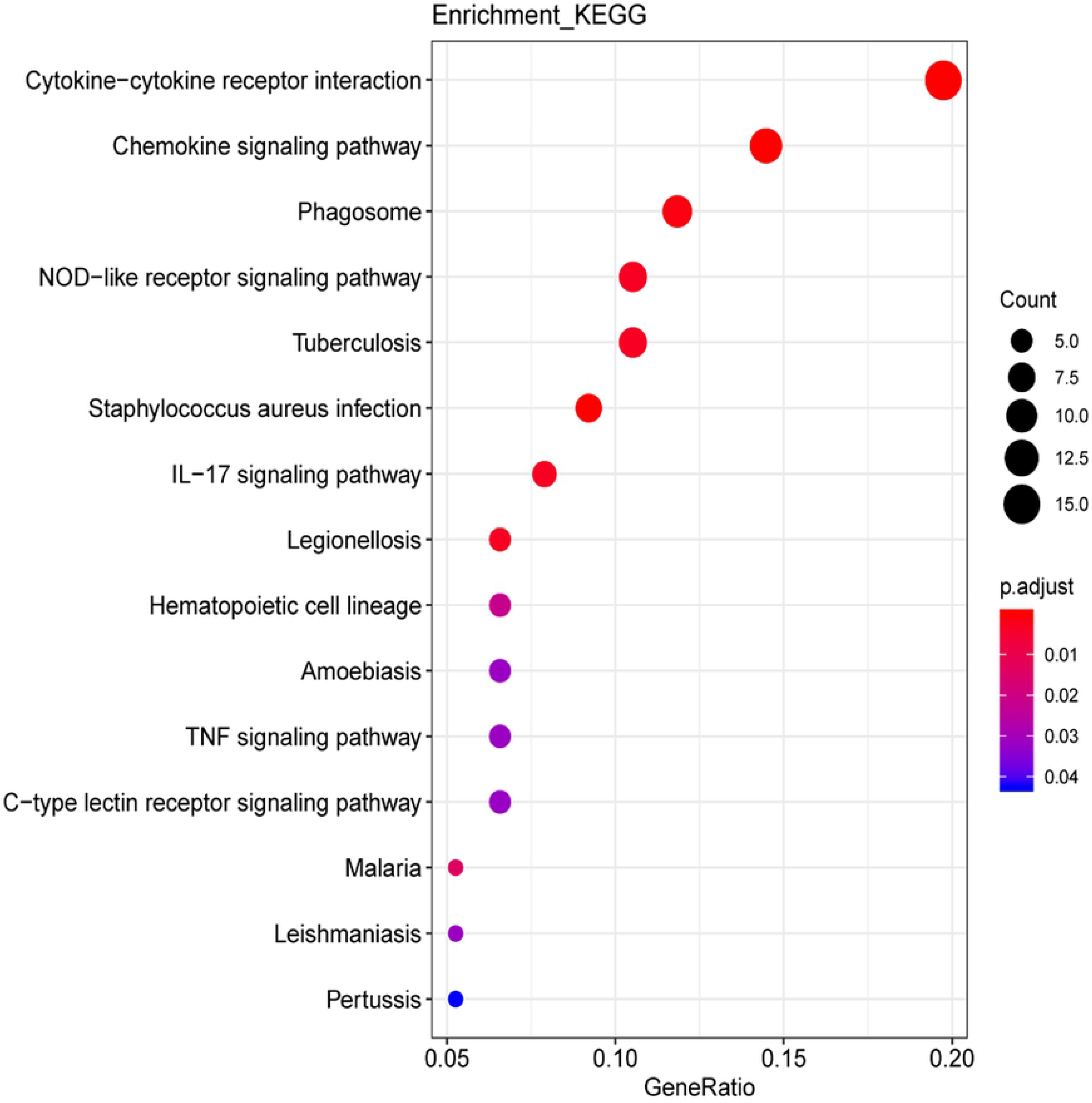
Top 15 enriched KEGG pathways analysis of significantly differentially expressed genes identified in mice with ankylosing spondylitis. The DEGs identified in mice with ankylosing spondylitis were significantly enriched in the Chemokine signaling pathway, NOD−like receptor signaling pathway, IL−17 signaling pathway and TNF signaling pathway.

**Figure. 3.**
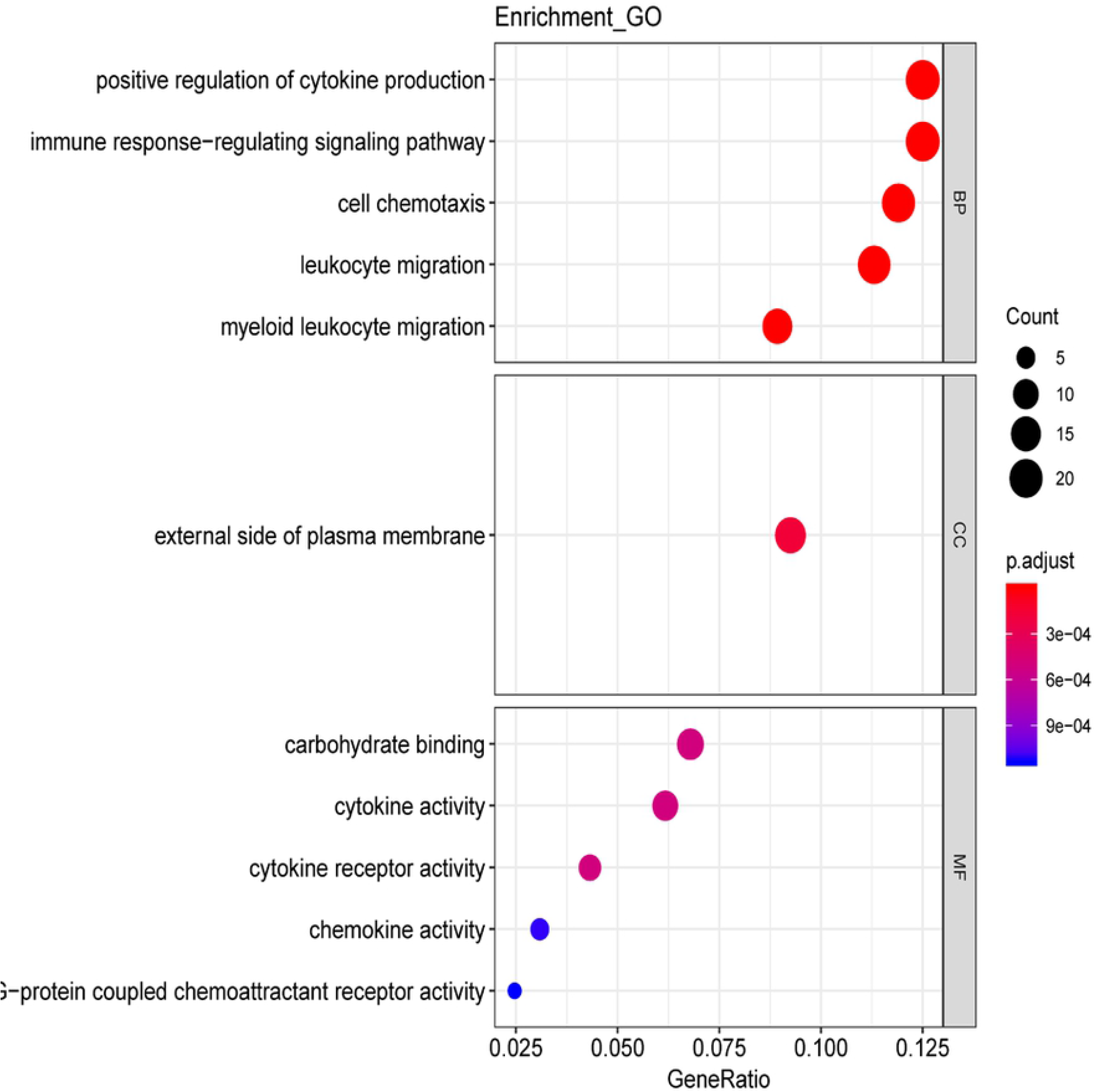
Top 11 most enriched gene ontology terms for the differentially expressed genes identified in mice with ankylosing spondylitis. BP: Biological Process GO-terms, CC: Cellular Component GO-terms, MF: Molecular Function GO-terms.

### 3.3. PPI network construction and hub genes analysis

When 230 specific DEGs were submitted to the STRING database, 128 genes were eliminated. The remaining DEGs were mapped into construct the PPI network (102 nodes and 372 edges) using the Cytoscape software (Fig. 4A). In the PPI network, the hub genes were sorted by the nodes number. In this study, twenty genes with highest degree scores were proposed as hub genes (Table 1). *MS4A6D* (degree=28), CTSS (degree=26), *FCGR1* (degree=25), *CD14* (degree=23), and *GPR65* had higher degrees in the PGISp mice, and were mainly associated with the NOD-like receptor, IL-17, and TNF signalling pathways. These core genes may play key regulatory roles in the pathogenesis of axSpA disease. In addition, sub-network clustering analyses were performed using the CytoHubba plugin, and two DEG-centred sub-networks were identified (Fig. 4B,C).

**Figure. 4.**
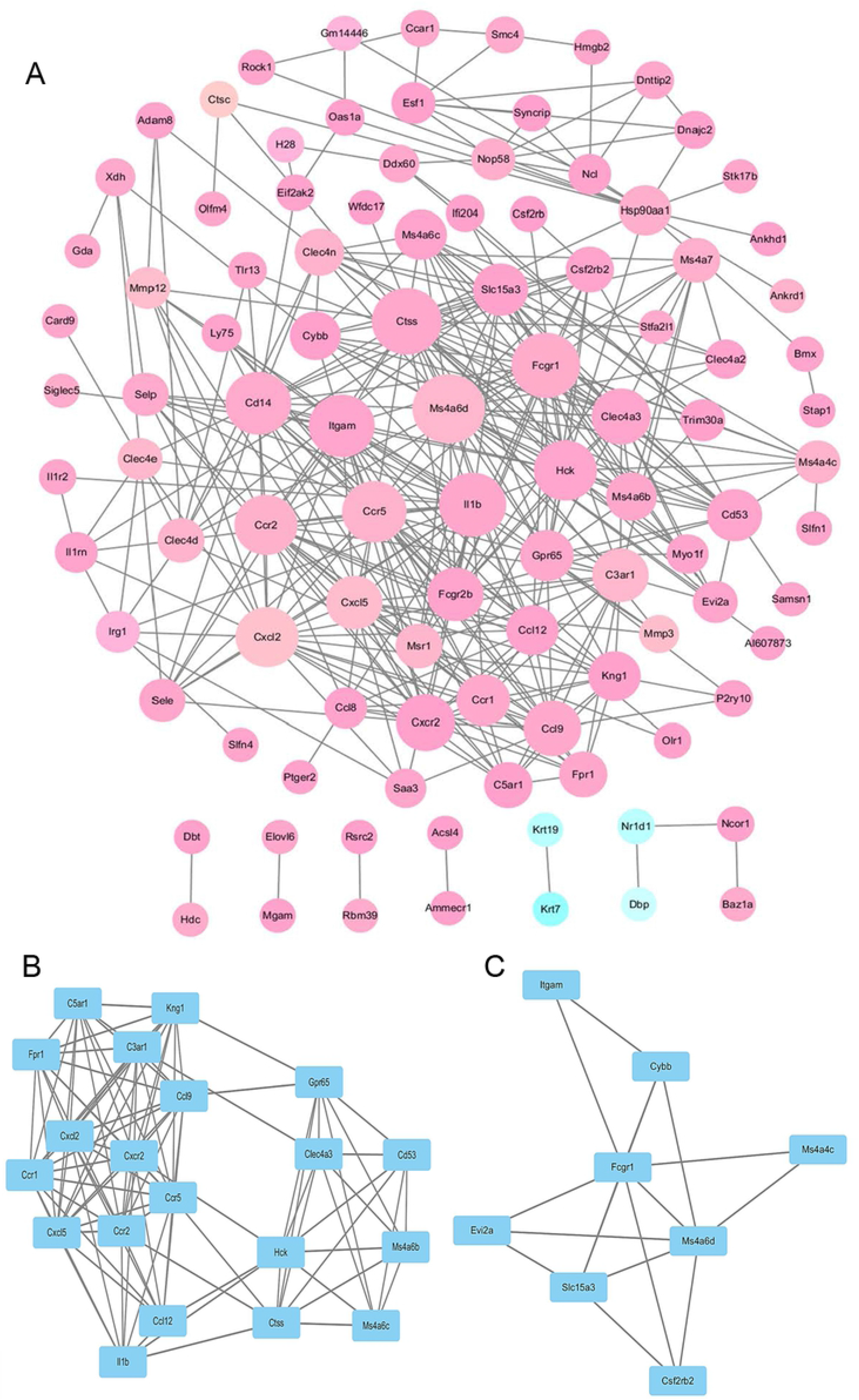
(A) Protein-protein interaction network for the differentially expressed genes identified in mice with ankylosing spondylitis. (B) and (C) Sub-networksof differentially expressed genes extracted from theprotein-protein interaction network. Nodes representthe proteins encoded byDEGs. The radius of the nods indicates the significance of enrichment, a pink color indicates that the DEG is upregulated and the green color indicates that the DEG is downregulated.

### 3.4. Analyses and construction of the TFs network

The the DEGs-related TFs regulatory network was identified with Cystoscope and is presented in Figure 5. A total of 10 TFs including *AP1, LSCBP, IRF1, RUNX1*, and *STAT3* and 48 DEGs were involved in the TF regulatory network. Several targeted regulatory genes overlapped between the TF series. The TF-DEG regulatory network consisted of 58 nodes and 77 lines. These TFs exhibited the ability to regulate DEGs which play vital roles in inflammatory response, immune response, and the TNF signalling pathway.

**Figure. 5.**
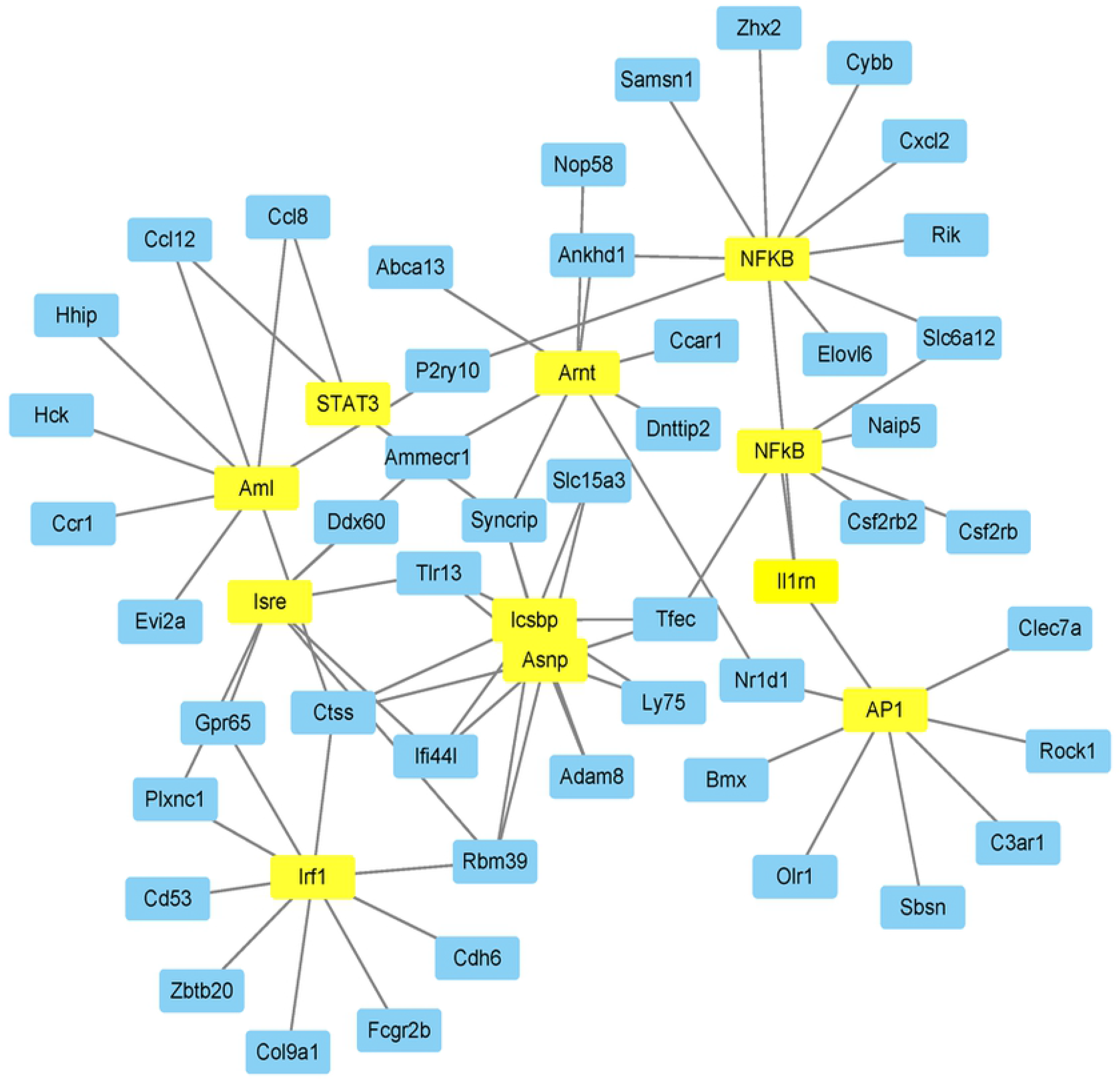
The transcriptional regulatory network.The yellow nodes represent TFs and the circle blue nodes represent the target genes.

### 3.5 Analysis and construction of microRNAs-target regulatory networks

A total of 10 significant microRNAs were screened by the WebGestalt database. Then the microRNAs-target regulatory networks were constructed, as shown in Figure 6. The microRNAs-target regulatory network contained 10 microRNAs, 26 target-specific genes, and 39 edges. MiR-142, miR-518a-2, miR-23, miR-452, miR-509, miR-365, and miR-150 were predicted to regulate the identified DEGs. These microRNAs regulate the inflammatory response, immune response, reactive oxygen species production, bone biology, and fibrosis, all of which play critical roles in the pathological process of axSpA.

**Figure. 6.**
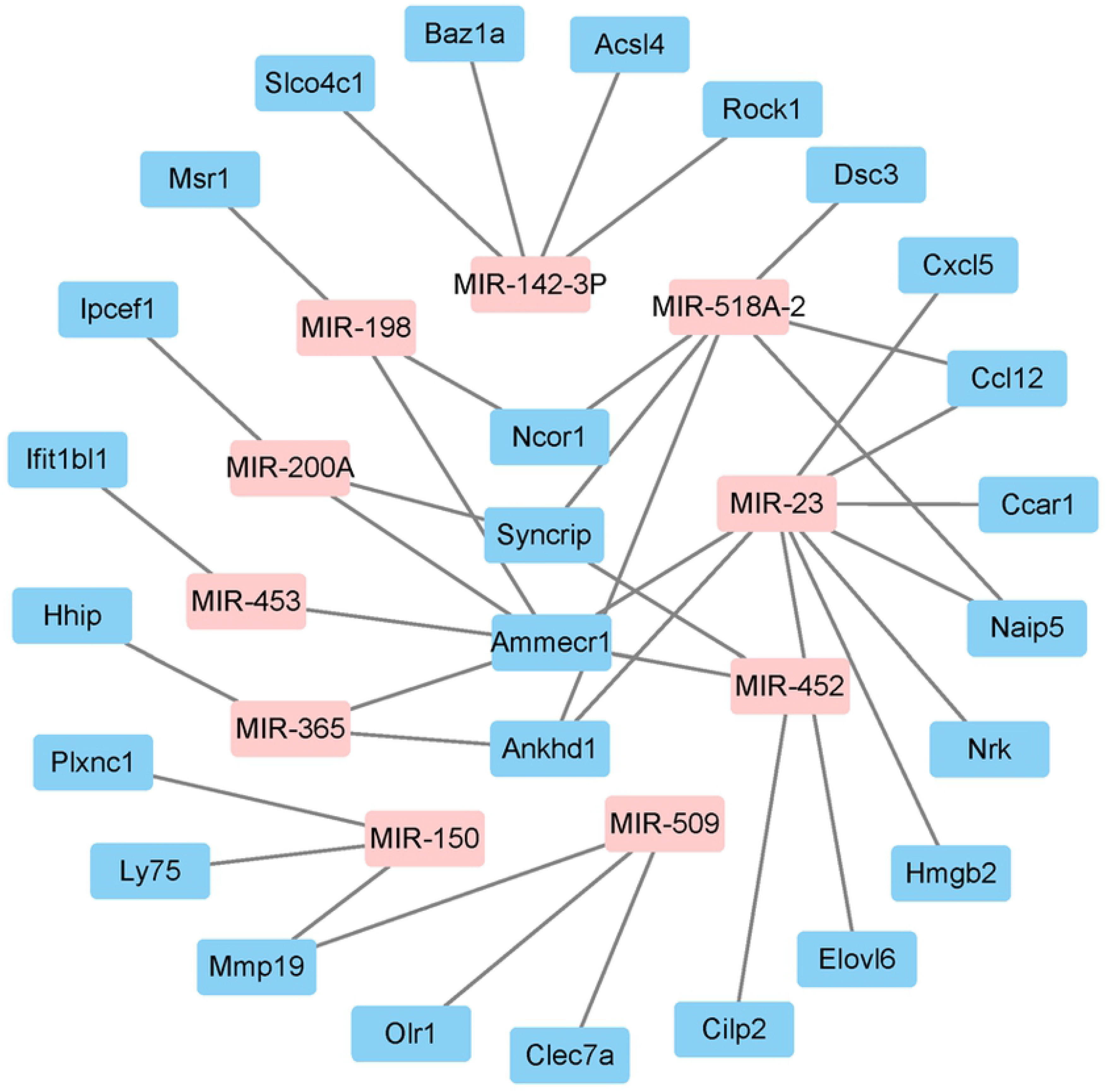
The microRNAs-target regulatory network.The pink nodes represent microRNAs and the circle blue nodes represent the target genes.

### 3.6. QPCR and histology validation

To testify the results of bioinformatics analysis, 9 hub gene, 3 microRNAs and 3 TFs were selected for RT-PCR and IHC respectively. Our findings indicated that the expression levels of the 7 hub genes, 2 microRNAs and 3 TFs were in accordance with the microarray data. The PG-induced mice had extensive chondrocyte proliferation and widely ankylosis of the sacroiliac joint compared with the control group (fig. 7).

**Figure. 7.**
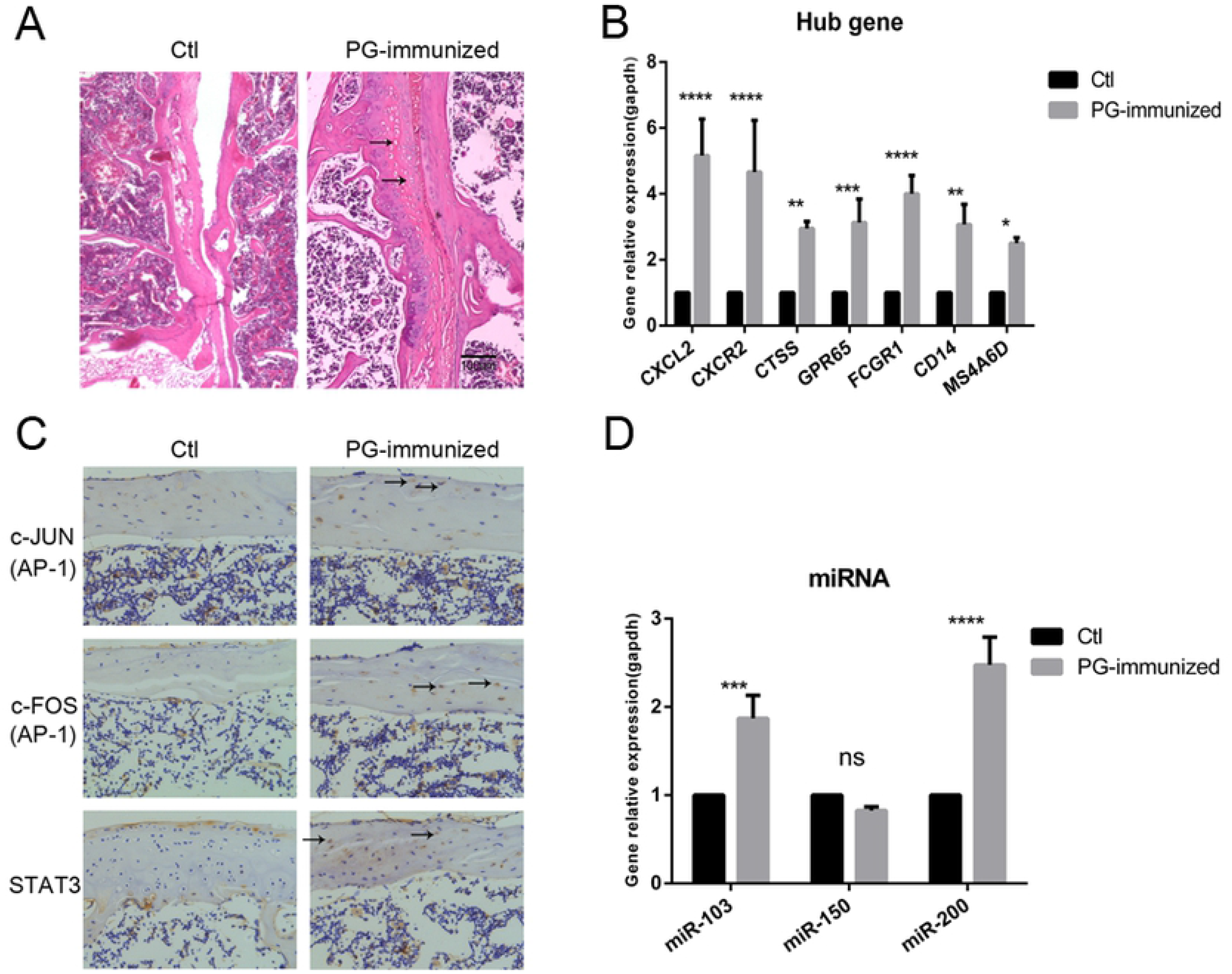
The relative expression of downstream hub genes, TFs and microRNAs in PG-induced mice and normal mice. (A) There were a mass of proliferation of chondrocyte-like cells (black arrow heads) in PG-induced sacroiliac joint, which formed a junction of chondrophytes, resulting in complete ankylosis of the sacroiliac joint (n=10).(B, D) The expression variations of hub genes and microRNAs between PG-induced mice and normal control. (C) Expression of transcription factors were determined by immunohistochemical staining of the sacroiliac joint from each group (n=10). \**p<0.05, **p<0.01, ***p<0.001, ****p<0.0001. ns=no significance.*

## 4. Discussion

In order to better elucidate the complex molecular mechanisms of axSpA, we performed in-depth analyses of microarray data (GSE13782) from PGISp mice and control mice. Through functional analyses and exploration, we identified 230 DEGs between the two groups, including 36 downregulated genes and 194 upregulated genes. Following this, the PPI network was constructed and two sub-networks were revealed involving *CXCL, CXCR, CTSS, GPR65* and *FCGR*. Furthermore, hub genes including *MS4A6D, CTSS, FCGR1*, and *CD14* were identified according to the degree counts. In addition, the TF regulatory network and microRNAs-target regulatory network were abstracted, which included 10 TFs and 10 microRNAs.

Previous studies have found that the chemokine, NOD-like receptor, IL-17, and TNF signalling pathways play crucial roles in the pathological process of axSpA(1, 19, 20), which is consistent with our analyses. Activation of the IL-23/IL-17 signal axis and secretion of pro-inflammatory cytokines lead to enthesitis, osteitis, and local joint inflammation (2). The role of NOD2 signalling in the induction of the IL-23/IL-17 axis is crucial for the promotion of local joint inflammation (21). Overexpression of TNF-α in mice reportedly generates enthesitis, aggressive polyarthritis, new bone formation and spinal arthritis (1). From the results of the sub-pathway clustering analyses and hub genes, several potential regulatory genes were identified. A previous study revealed that *FCGR2B* gene polymorphisms are significantly associated with autoimmune diseases, including axSpA (22), particularly in autoimmunity and osteoclast differentiation. Furthermore, the results of the study by Xu and this study identified the upregulation of *FCGR1B*, which suggests that its expression may be related to the pathological process of axSpA. Meanwhile, FCGR is a major candidate gene in autoimmune disease pathogenesis; it reduces the expression and function of inhibitory FcγRII on macrophages and activated B cells (23). Wang (24)and Lee (25) both identified a significant association between FGCR and axSpA. Spondyloarthritis patients showed an increased expression of G-protein coupled receptor (*GPR65*) compared to healthy controls(26), which was coincident with our findings. Wirasinha (27) also demonstrated that GPR65 exacerbates autoimmune disease through CD4+ T cells and invariant NK T cells. Therefore, finding a small-molecule inhibitor of GPR65 may be a promising therapeutic approach for axSpA. Furthermore, previous studies all showed that CXCL, CXCR, CCL, CCR, CYBB, ITGAM, and IL-1β are involved in regulating the NOD-like receptor, IL, and TNF signalling pathways ((28–31).

Based on the TF-DEG network constructed in this study, two TFs, STAT3, and LSCBP, were identified as particularly important in axSpA. *STAT3* is a gene involved in the IL23/Th17 signalling pathway in inflammatory diseases, including subclinical Crohn’s-like colitis (32). Decreases in p-STAT3 could inhibit the expression of ARG1 and T cell suppressive function in myeloid-derived suppressor cells in patients with axSpA (33). The secretion of IL-17A could drive osteoblast differentiation by promoting phosphorylation of JAK2 and STAT3 (34). Studies have revealed that STAT3 is involved in IL-17A/IL23-induced axSpA, and acts as a key TF of osteoblast differentiation. Choi (35) confirmed that LSCBP (IRF8) binds to the Spp1 promoter region to repress the immune response and OPN expression, which is involved in bone remodelling in axSpA (35). NFKB was also identified as a candidate gene in axSpA in a previous study (36), consistent with our analyses.

From the microRNAs-DEG network, we determined that many microRNAs and pathways are closely associated with axSpA. MiR-150-5p increases the production of inflammatory cytokines and the function of dendritic cells by promoting the nuclear translocation of c-fos, NF-kB p65, and the JAK/STAT pathway (37). MiR-150-5p also promotes osteoblast function and bone mineralisation by targeting MMP14 in axSpA (38). The upregulation of miR-200 drives a partial inhibition of the MYD88-dependant signalling branch and provokes TFs associated with the TICAM-dependent signalling branch in systemic lupus erythematosus (SLE) (39). Furthermore, several recent studies have shown that miR142, miR23, miR452, miR509, and miR365 are closely associated with the immune response, inflammatory response, and bone metabolism. Therefore, further investigations will be beneficial in examining these microRNAs with respect to axSpA.

## 5. Conclusion

In conclusion, we analysed the gene expression profiles of five PGISp mice and three naïve controls. In this study, a total of 230 candidate genes were identified, including 36 downregulated genes and 194 upregulated genes. The results indicated that the chemokine, NOD-like receptor, IL-17, and TNF signalling pathways, as well as genes associated with these pathways, are associated with the pathogenesis of axSpA. STAT3 and LSCBP were identified as particularly important TFs in axSpA through regulating inflammatory response and osteoblast differentiation. MiR-150-5p and miR-200 are potential microRNAs involved in the signalling branches of bone metabolism and immune response. Finally, we validated the findings of bioinformatics analysis with QPCR and histology staining. Our finding provides insights into the pathomechanisms of axSpA, as well as potential research directions. Further experiments are warranted to validate these findings and to explore their roles in the pathogenesis of axSpA.

## Authors’ contributions

G. Liu and S. Liang designed this study; Zhen-zhen Zhang performed the experiments; Y. Zou and J. Zeng analyzed the data; Hai-hong Li drafted the manuscript.

